# Target of selective auditory attention can be robustly followed with MEG

**DOI:** 10.1101/588491

**Authors:** Dovilė Kurmanavičiūtė, Hanna Kataja, Mainak Jas, Anne Välilä, Lauri Parkkonen

**Affiliations:** Department of Neuroscience and Biomedical Engineering, Aalto University, P.O. Box 12200, FI-00076 Aalto, Finland; Aalto NeuroImaging, Aalto University, FI-00076 Aalto, Finland; Athinoula A. Martinos Center for Biomedical Imaging, 149 Thirteenth Street, Suite 2301, Charlestown, Massachusetts 02129, USA

## Abstract

Selective auditory attention enables filtering of relevant acoustic information from irrelevant. Specific auditory responses, measurable by magneto- and electroencephalography (MEG/EEG), are known to be modulated by attention to the evoking stimuli. However, such attention effects have typically been studied in unnatural conditions (e.g. during dichotic listening of pure tones) and have been demonstrated mostly in averaged auditory evoked responses.

To test how reliably we can detect the attention target from unaveraged brain responses, we recorded MEG data from 15 healthy subjects that were presented with two human speakers uttering continuously the words “Yes” and “No” in an interleaved manner. The subjects were asked to attend to one speaker. To investigate which temporal and spatial aspects of the responses carry the most information about the target of auditory attention, we performed spatially and temporally resolved classification of the unaveraged MEG responses using a support vector machine.

Sensor-level decoding of the responses to attended vs. unattended words resulted in a mean accuracy of 79% ± 2% (*N* = 14) for both stimulus words. The discriminating information was mostly available 200–400 ms after the stimulus onset. Spatially-resolved source-level decoding indicated that the most informative sources were in the auditory cortices, in both the left and right hemisphere.

Our result corroborates attention modulation of auditory evoked responses and shows that such modulations are detectable in unaveraged MEG responses at high accuracy, which could be exploited e.g. in an intuitive brain–computer interface.

## Introduction

Selective auditory attention enables filtering of relevant acoustic information from irrelevant and is often studied using dichotic listening^1, 2^ where the listener is exposed to simultaneous but different auditory streams to each ear and is asked to follow one stream while suppressing the other, akin to the cocktail party problem^3^. Selectively attending to one stream manifests as changes in auditory evoked responses that can be measured non-invasively with electroencephalography (EEG) and magnetoencephalography (MEG)^4–8^.

More recently, machine-learning methods have been applied to EEG/MEG data to study attention modulation of transient auditory evoked responses^9^, auditory steady-state responses^10, 11^ or responses to continuous speech^12–15^. Exploiting such attention modulation in a brain–computer interface has been probed in several studies^16–25^, some of which have employed natural sounds as stimuli and yielded a useful-in-practice classification accuracy also when applied to patients that cannot communicate^26, 27^. However, these auditory speller -type BCI systems require extensive training that might be exhausting for a patient. Furthermore, in patients with disorders of consciousness, using this type of a BCI may exceed the capacity of their working memory^25, 28^, which could drastically drop the accuracy. In comparison to a speller-BCI, BCIs based on either speech tracking or P300 auditory streaming could be designed such that their working-memory load is limited^21, 29–31^. However, a BCI utilizing speech tracking requires long data spans (usually tens of seconds) to output one bit whereas auditory streaming BCIs can operate on short trials (one or few seconds).

Auditory streaming BCIs often employ oddball streams, comprising frequently-occurring stimuli (standard) and a rarely-occurring exception (deviant). Selective attention then increases the amplitude of the response to a deviant compared to an unattended deviant. However, this approach allows attention target to be determined only at the rate the deviants are presented, and this rate cannot be increased above 10 or 20% of all stimuli without diminishing the overall amplitude of the deviant responses. Therefore, the bitrate of such a BCI remains modest.

In this study, we propose a novel, naturalistic paradigm that enables more rapid tracking of the target of attention while also providing a task allowing behavioural quantification of the deployment of attention. To this end, we created an acoustically realistic scene with two concurrent auditory stimulus streams. Stimuli comprised of two human speakers uttering the words “yes” and “no” in an interleaved manner at −40 and +40 degrees from the line forward from the subject, mimicking a real-life situation where two persons are speaking simultaneously on the sides of the subject. In each stream, the pitch of the word alternated (standard) but this implicit rule was occasionally broken by presenting two same-pitch versions of the stimulus word in succession (deviant). We measured MEG in 15 subjects while they were presented with these stimuli and were asked to covertly count these deviants in the attended stream and report the number at the end of each measurement block.

## Results

### Behavioral data

On average, the subjects reported 40±17.6% (deviant probability 10%, *N* = 5) and 97 ±0% (deviant probability 5%, *N* = 6) of the deviants in the stream they were instructed to attend to. Three subjects were not included in this analysis due to technical problems in collecting their deviant counts.

### Sensor-level analysis

Average evoked responses to each attention condition (“Attended Yes”, “Unattended Yes”, “Attended No”, “Unattended No”) are shown for a single subject in Fig. 1 and for the group in Fig. 2. Each condition represents pooled responses to the low- and high-pitch stimuli. It is important to note that these average evoked responses were computed only for the sensor- and source-level visualizations, and all decoding was performed on unaveraged (single-trial) responses.

**Figure 1.**
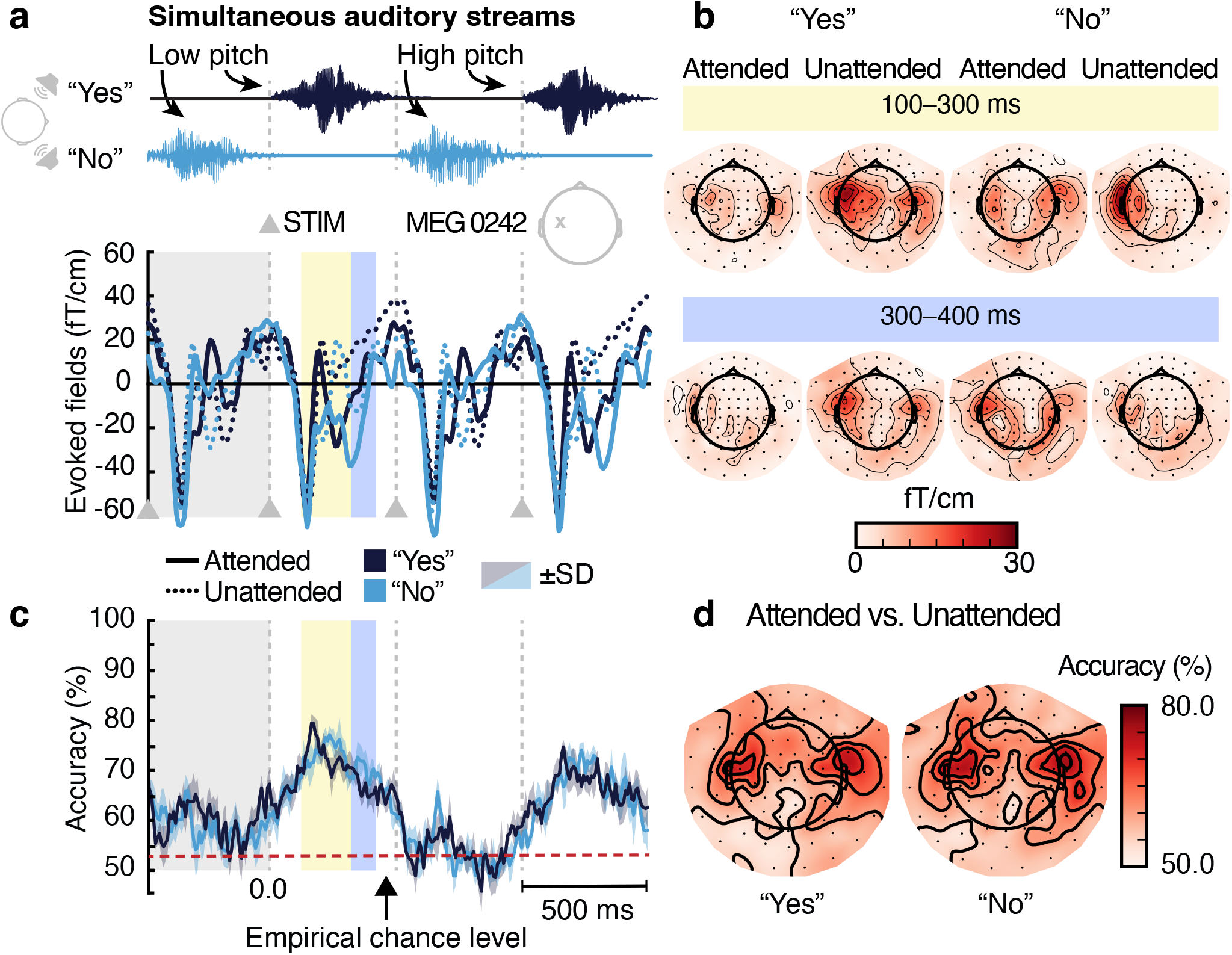
Evoked responses and decoding accuracy in a representative subject (S03) to four consecutive stimuli in a 2-s window. **a**: Acoustic waveform of the stimuli and the corresponding average responses at a planar gradiometer channel low-pass-filtered at 30 Hz in the attended (solid lines) and unattended (dotted lines) conditions. **b**: Spatial patterns (gradient strength maps showing the root mean square for each gradiometer pair) of the evoked responses. **c**: Time-resolved decoding of attended vs. unattended “Yes” (dark blue) and attended vs. unattended “No” (light blue) mean accuracy and standard deviation (dark blue and light blue shading) computed across the cross-validation folds. **d**: Spatially-resolved decoding accuracy at the sensor level for the same attention conditions.

**Figure 2.**
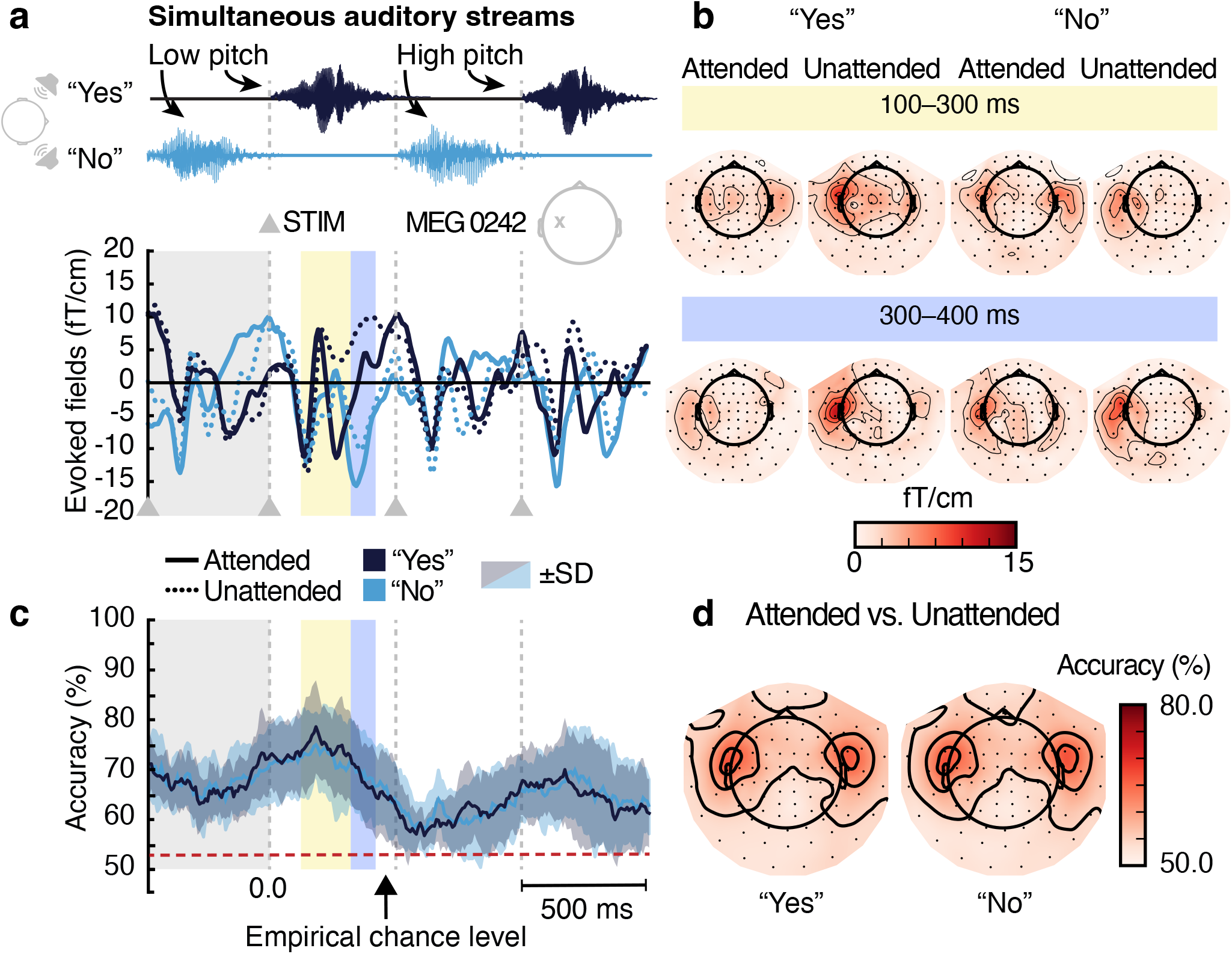
Evoked responses and decoding accuracy averaged across the group (*N* = 14) to four consecutive stimuli in a 2-s window. **a**: Acoustic waveform of the stimuli and the corresponding average responses at a planar gradiometer channel low-pass-filtered at 30 Hz in the attended (solid lines) and unattended (dotted lines) conditions. **b**: Spatial patterns (gradient strength maps, showing the root mean square for each gradiometer pair) of the evoked responses. **c**: Time-resolved decoding of attended vs. unattended “Yes” (dark blue) and attended vs. unattended “No” (light blue) mean accuracy and standard deviation (dark blue and light blue shading) computed across the cross-validation folds. **d**: Spatially-resolved decoding accuracy at the sensor level for the same attention conditions.

The group-averaged evoked responses peaked at 250 ms (at channel ‘MEG 1322’) after the stimulus onset for the “Attended Yes” and at 136 ms (‘MEG 1322’) for “Unattended Yes” condition. For the condition “Attended No”, responses peaked at 340 ms (‘MEG 0242’) and for “Unattended No” at 350 ms (‘MEG 0212’). The planar gradient strength maps (Fig. 1a and Fig. 2b) are compatible with sources in auditory cortices.

Time-resolved decoding was performed on the unaveraged epochs comprising all channels at each time point. At the group level, decoding “Attended No” vs. “Unattended No” and “Attended Yes” vs. “Unattended Yes” both showed peaks around 160 ms (Fig. 2c).

Spatially-resolved decoding indicated that the most informative signals arose from temporal regions; the patterns of decoding accuracy were qualitatively similar across the subjects; see Fig. 1d for a representative subject and Fig. 2d for the group result.

To aim at the highest accuracy in determining the direction of attention, we also decoded using the entire epoch, that is, all time points and all channels at once. First, we tested with 1-s epochs (–200… 800 ms) which yielded a mean accuracy of 79%±2% (range 67–91%) for “Attended No” vs. “Unattended No” and 79%±2% (range 68–91%) for “Attended Yes” vs. “Unattended Yes”; see Fig. 3. The group mean accuracy was not significantly different between the stimulus words (paired t-test; *p* = 0.874).

**Figure 3.**
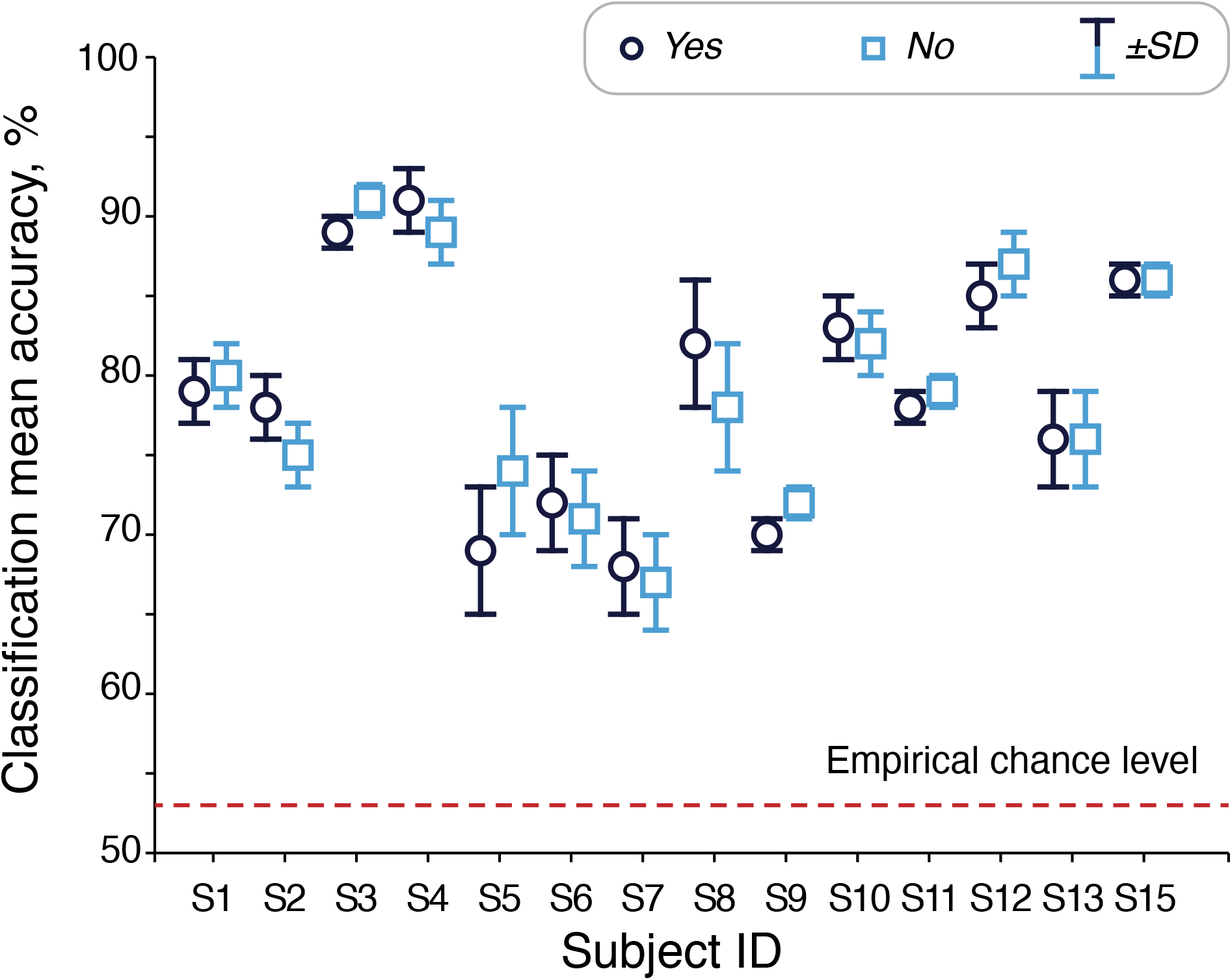
Decoding accuracy of responses to attended vs. unattended “Yes” and “No” word-stimuli for all subjects using the 1-s epoch. SD is shown as plot whiskers and indicates standard deviation across the cross-validation folds for each subject.

Prolonging the decoding epoch to 2 s (–500… 1500 ms) resulted in an average decoding accuracy of 83%±2% (range 71–94%) for “Attended Yes” vs. “Unattended Yes” and 83%±2%; (range 68–94%) for “Attended No” vs. “Unattended No”. Using the 2-s vs. 1-s epochs increased the classification accuracy in all 14 subjects, which is a significant change (binomial test: p =0.000061).

To further characterize the stimulus-related information present in the auditory evoked responses, we decoded also for the stimulus word (not for attention). The mean accuracy was 88%±2% (range 75–96%) for the 1-s epochs and 93%±1% (range 82–99%) for the 2-s epochs when decoding “Attended Yes” vs. “Attended No”. Similarly, when decoding for “Unattended Yes” vs. “Unattended No”, we obtained an average decoding accuracy of 88%±2% (range 76–95%) for 1-s epochs and 92%±2% (range 77–98%) for the 2-s epochs. Thus, the accuracy of word-wise decoding did not depend significantly on whether the words were attended or not (p =0.80 for the short epochs and p =0.53 for the long epochs). Compared to attention decoding, this word-wise decoding gave statistically significantly higher accuracy for both the short (p < 0.005 for all four possible comparisons) and long (p < 0.005) epochs.

We also performed pitch-wise decoding (unaveraged evoked responses to high-pitch vs. low-pitch versions of the word stimuli), which yielded above chance-level decoding accuracy of 70%±3% (“Attended No”), 71%±3% (“Unattended No”), 71%±3% (“Attended Yes”) and 72%±3% (“Unattended Yes”) across all subjects (N = 14).

### Source-level analysis

The group-level source estimates depicted in the Fig. 4 show the responses to attended and unattended word stimuli at three different latencies. In the right hemisphere, the activation peaked at 270 ms after the onset of the attended “Yes”. Activation to the attended “No” peaked at 330 ms in the left hemisphere after the stimulus onset.

**Figure 4.**
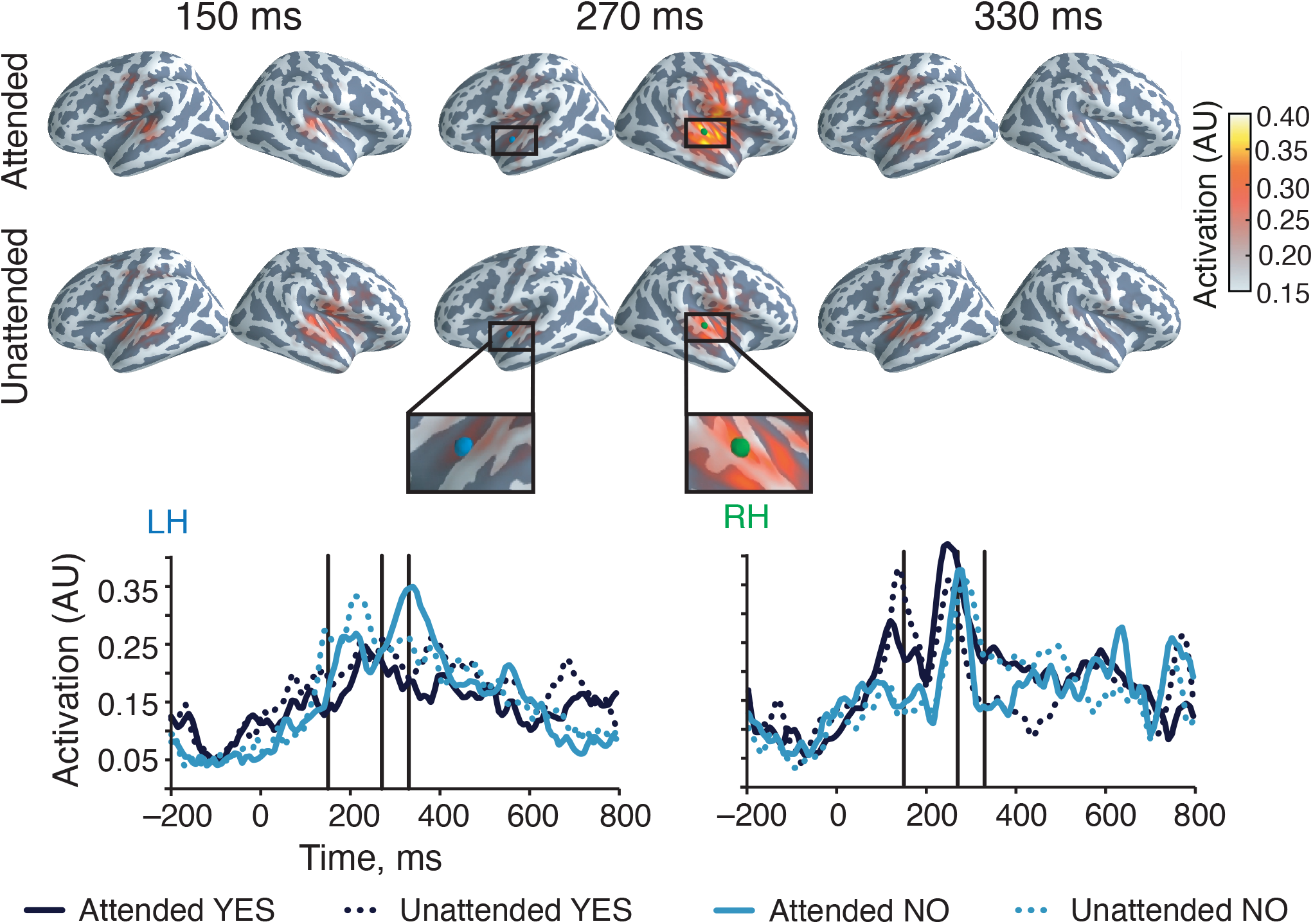
**Top**: Group-level source estimates of the responses to attended and unattended word stimuli at three latencies after the stimulus onset; *N* = 11. The color represents the source amplitude first normalized to the absolute peak value of each individual source estimate and then averaged across subjects. **Bottom**: Left (LH) and right (RH) auditory-cortex activation to both stimulus words (“Yes”/”No”) and attention conditions (“attended”/”unattended”) extracted from the source estimate at the colored dots (green/blue).

Spatial-searchlight decoding (Fig. 5) revealed that the source signals giving the highest decoding accuracy arose from the auditory cortices and sensorimotor cortex that has been associated with auditory stimuli processing as well. However, the spatial peaks of accuracy, as show in Fig. 5, did not align well in time and space across subjects, which led to low group-average accuracy for any single location on the cortex.

**Figure 5.**
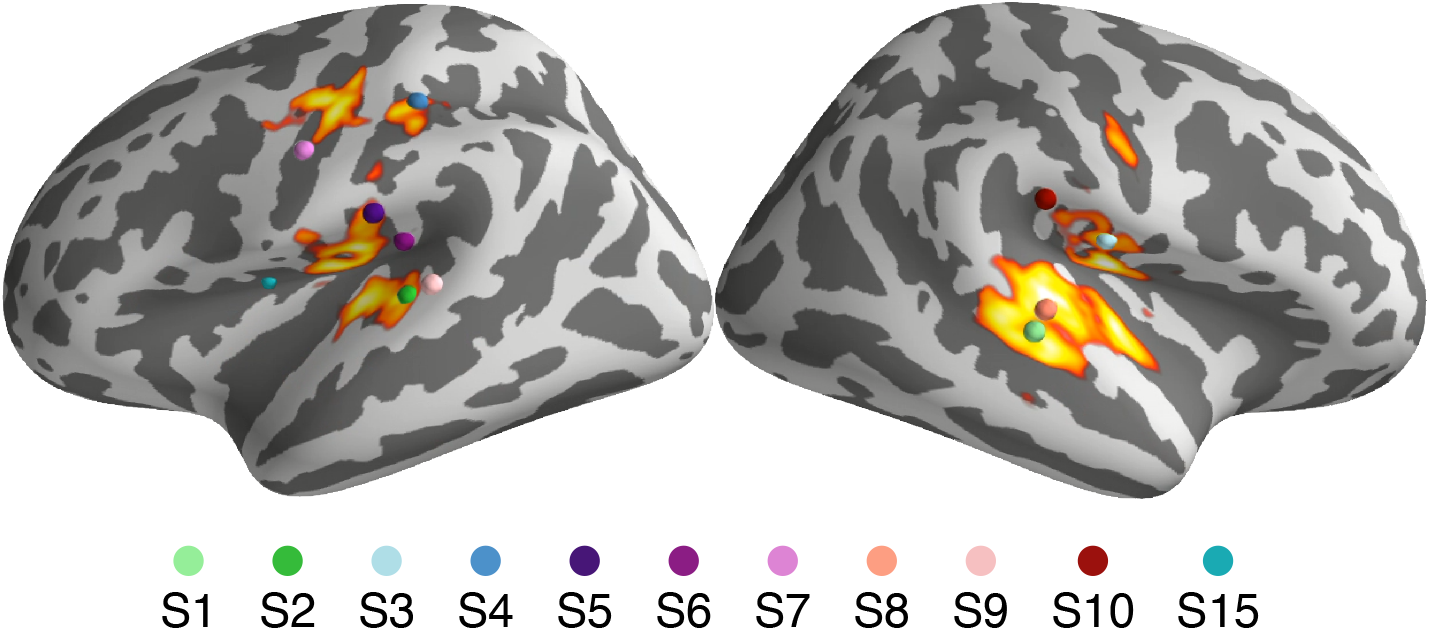
Spatial-searchlight decoding accuracy peaks in all subjects (N =11; each color dot represents the peak in one subject). in classification of attended vs unattended word (“Yes” and “No” averaged together).

## Discussion

In this study, we recorded brain signals during a “cocktail-party scene” and demonstrated that the target of selective auditory attention to concurrent streams of naturalistic spoken-word stimuli can robustly (accuracy 79%) be decoded from just 1 s of MEG data. The decoding accuracy peaked around 160 ms after the onset of both stimulus words and remained above chance level for several hundred milliseconds. The highest accuracy was obtained with signals arising from the auditory cortices.

Previous studies have shown that non-semantic acoustic properties, such as sound-source-specific pitch^32^, are crucial for solving the cocktail party problem at the perceptual level^33, 34^. The cortical representation of these properties and the MEG signals evoked by them may contribute to our ability to decode the target of attention. The presence of pitch-related information in our data was demonstrated by the above-chance-level accuracy when decoding for the pitch (low vs. high pitch) instead of attention. Similarly, we obtained a high accuracy when decoding for the stimulus word (group mean of 88% for 1-s epochs) or for the stimulus pitch (range 70– –72%) instead of attention, which also speaks for the presence acoustic information in the MEG responses; differing stimulus duration is probably the most important discerning factor.

Earlier studies have demonstrated that rich naturalistic stimuli, compared to monotonous tones, not only improve the users’ ergonomic evaluation of the situation but also yield higher decoding accuracy^21, 35^. Further, it has been shown that subjects perform better on selective attention tasks when presented with naturalistic speech in comparison to other kind of naturalistic stimuli^36^.

In our study, the pitch difference between the spoken-word streams due to speaker gender (male and female voices) likely helped focusing attention to one speaker. Yet, the pitch itself did not seem to play a role in either stream alone as the attention decoding accuracy for each word-stream was very similar (Fig. 3).

Stimulus timing likely has an effect on decoding accuracy as it influences the amplitude and latency of attention-modulated evoked responses. Stimulus onset asynchrony (SOA) has been shown to affect accuracy in decoding attention to simple tones by Höhne and colleagues^37^; they found that SOA of 1000 ms gave the best decoding accuracy but the highest information transfer rate was achieved with short SOA’s (87–175 ms). Other studies that used virtual sound stimuli observed that SOA of 400–600 ms provided the best decoding accuracy^28, 38^. Given those previous studies, our SOA of 1100 ms was likely close to optimal in terms of decoding accuracy but probably would not yield the highest information transfer rate, if our paradigm was applied in a brain–computer interface (BCI).

Typically, using a longer span of data for decoding improves accuracy if all data are informative. Also our results showed that using the long (2-s) instead of the short (1-s) epoch increased the accuracy of decoding the target of attention in all 14 subjects. Again, for a BCI, the long epochs may not be the optimal choice to maximize the bitrate.

Spatial searchlight decoding across the cortex yielded accuracy peaks at locations similar to those of the largest differences in the source estimates of the two attention conditions. This agreement of the two analysis methods further supports the notion that the selective auditory attention-modulated cortical activity is mostly in the primary auditory cortex^39^. Regardless of the roughly similar cortical location of the most attention-informative source in each subject, these locations did not fully overlap, which led to a dispersed group average even though interindividual variation of cortical anatomy was reduced by surface-based morphing of the individual brains to an average brain. This variation – although minor – in the location and orientation of the source providing the highest decoding accuracy likely means that classifiers do not generalize well across subjects but that the classifier should be trained separately for each subject if one is aiming to the highest accuracy.

It is conceivable that in our experiment participants applied different strategies of keeping their attention to one auditory stream or they might have even changed their strategy in the course of the experiment. This possibility could be studied by training the decoder by samples from specific parts of the recording (e.g. only from the beginning) and comparing the obtained classification accuracy. In addition, the influence of stimulation rate to selective attention and its decoding from brain signals could be tested. Moreover, future studies could assess individual differences in response latency and spatial patterns on the MEG sensor array that may limit across-subject generalization.

Using the current experimental paradigm, one could test how robustly the observed attention modulation of brain responses could be detected by EEG instead of MEG. Spatial separability of cortical sources is typically poorer in EEG compared to MEG and thus the accuracy of decoding the attended word stream from EEG would likely be lower; yet, the accuracy could remain at a level which enables an intuitive auditory-streaming brain–computer interface.

## Conclusions

We showed that the attended spoken-word stream can reliably be decoded from just one-second epochs of unaveraged MEG data. The achieved high decoding accuracy shall enable future investigations on the neural mechanisms of attentional selection and it may also be exploited in a MEG- or EEG-based streaming brain–computer interface.

## Materials and methods

### Participants

Fifteen healthy adult volunteers (4 females, 11 males; mean age 28.8±3.8 years, range 23–38 years) participated in our study. Two subjects were left-handed and the rest right-handed. Participants did not report hearing problems or history of psychiatric disorders. The study was approved by the Aalto University Research Ethics Committee. The research was carried out in accordance with the guidelines of the Declaration of Helsinki, and the subjects gave written informed consent prior the measurements.

### Stimuli and experimental protocol

The subjects were presented with two auditory streams, one comprising the spoken word “Yes” and the other the word “No”. The words alternated such that the words did not overlap. In each stream (“Yes” and “No”), the stimulus onset asynchrony (SOA) was 1100 ms, and the durations of the stimulus words were 450–550 ms depending on the word and its pitch variant. Fig. 6B illustrates the stimulus sequence.

**Figure 6.**
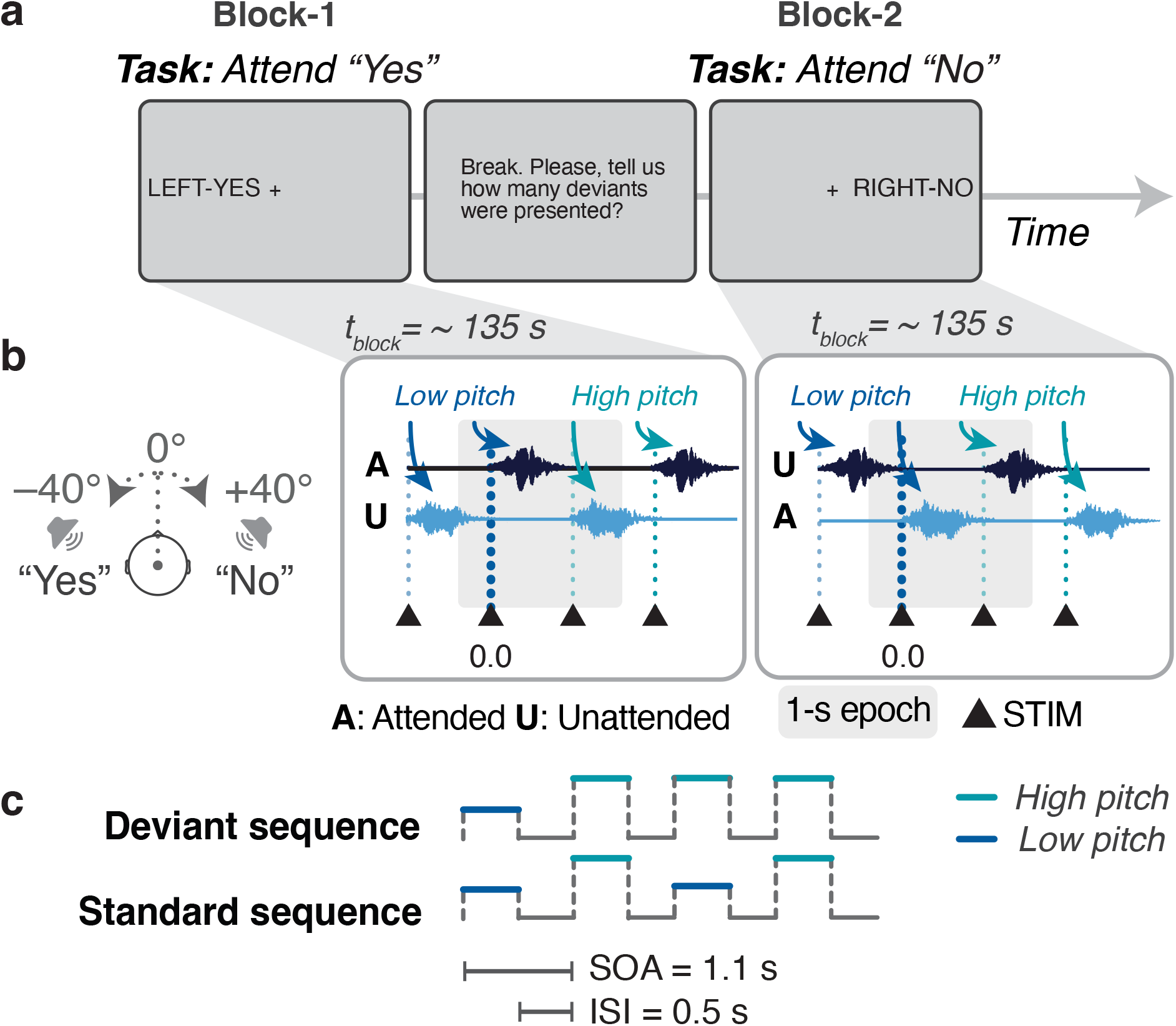
Experimental design. **a:**Block structure and the instructions to the subject. **b:**Extraction of epochs for decoding responses to attended vs unattended “Yes” and “No” words. Stimulus onsets are marked with triangles, colored according to pitch and attention. The grey shading indicates the short (1-s) epoch. **c:**Stimulus sequence excerpts with and without a deviant.

To create a realistic acoustic scene, the stimuli were recorded with a dummy head (Mk II, Cortex Instruments GmbH, Germany) at the center of a room with dimensions comparable to those of the magnetically shielded room where the MEG recordings were performed later. The speakers were standing at about at −40 and +40 degrees from the front-line of the dummy head at a distance of 1.13 m. The word “Yes” was uttered by a female and the word “No” by a male speaker. Thus, in the experiment, the sound of each speaker was presented to both ears of the subject but with different amplitudes and phases; see Fig. 6B.

Subjects were asked to attend to one of the two asynchronously alternating spoken word streams at a time by a visual instruction, see Fig. 6a. Each stream had two alternating pitches of spoken word (denoted as […, *yes, YES, yes, YES,*…] and as […, *no, NO, no, NO,*…]). The original voice recordings were used as the low-pitch stimuli, and the pitch was increased by 13% and 15% for the high-pitch versions of “Yes” and “No”, respectively. In each spoken-word stream, occasional violations (deviants) of otherwise regular pitch alternation (standards) occurred. The subjects were instructed to count the deviants in order to keep their attention to the indicated spoken word stream stimuli. The stimulus sequence always started with the low-pitch word; see Fig. 6b.

Deviants were presented with probability of 10% in both streams for the first seven subjects and with probability of 5% for the rest of the subjects. The deviant frequency was decreased based on subject feedback to reduce the mental load of memorizing the deviant count.

The experiment comprised 8 blocks, each lasting about 5 min. Two seconds before a block started, the subject was instructed to direct his/her attention to one of the streams by the cues “LEFT-YES” or “RIGHT-NO” on the screen. The task of the subject was to focus on the indicated word stream, covertly count the deviants, maintain gaze at the fixation cross displayed on the screen and verbally report the count at the end of the block.

During the cue “LEFT-YES”, the evoked responses to the word “Yes” were assigned to the condition “Attended Yes” and the evoked responses to “No” were assigned to the condition “Unattended No”. Similarly, during the cue “RIGHT-NO”, evoked responses to “No” were assigned to the condition “Attended No” and, accordingly, evoked responses to “Yes” were assigned to the condition “Unattended Yes”.

The experiment always started with a block with the cue “LEFT-YES” and was followed by a block with the cue “RIGHT-NO”. The order of the remaining six blocks was randomized across subjects. The first blocks were not randomised due to our main goal to use the first two blocks for training the classifier. The total length of the experiment was 50–60 minutes including the breaks between the blocks.

PsychoPy version 1.79.01^40, 41^ Python package was used for controlling and presenting the auditory stimuli and visual instructions. The stimulation was controlled by a computer running Windows 2003 for the first nine subjects and Linux Ubuntu 14.04 for the rest. Auditory stimuli were delivered by a professional audio card (E-MU 1616m PCIe, E-MU Systems, Scotts Valley, CA, USA), an audio power amplifier (LTO MACRO 830, Sekaku Electron Industry Co., Ltd, Taichung, Taiwan), and custom-built loudspeaker units outside of the shielded room and plastic tubes conveying the stimuli separately to the ears. Sound pressure was adjusted to a comfortable level for each subject individually. Due to timing inaccuracies in the stimulus presentation system, the delay from the trigger to sound onset for the “Yes” stimuli varied with a standard deviation of 7 ms while that for “No” varied with a standard deviation of 11 ms.

### MEG data acquisition

MEG measurements were performed with a whole-scalp 306-channel Elekta Neuromag VectorView MEG system (MEGIN Oy, Helsinki, Finland) at the MEG Core of Aalto Neuroimaging, Aalto University. During acquisition, the data were filtered to 0.1–330 Hz and sampled at 1 kHz. Prior to the MEG recording, anatomical landmarks (nasion, left and right preauricular points), head-position indicator coils, and additional scalp-surface points (around 100) were digitized using an Isotrak 3D digitizer (Polhemus Navigational Sciences, Colchester, VT, USA). Bipolar electrooculogram (EOG) with electrodes positioned around the right eye (laterally and below) was recorded. Fourteen of the 15 subjects were recorded with continuous head movement tracking. All subjects were measured in the seated position. The back-projection screen for delivering the visual instructions was 1 m from the eyes of the subject. If needed, vision was corrected by nonmagnetic goggles.

The MEG recording of one subject had technical problems and this dataset had to be dropped from the analysis.

### Data pre-processing

The MaxFilter software (version 2.2.10; MEGIN Oy, Helsinki, Finland) was applied to all MEG data (magnetometers and planar gradiometers) to suppress external interference using temporal signal space separation and to compensate for head movements^42^. Further analysis was performed using the MNE-Python [43, 44, *version 0.21*;] and ScikitLearn [45, *version 0.23.2*;] software packages.

Infinite-impulse-response filters (4^th^-order Butterworth, applied both forward and backward in time) were employed to filter the unaveraged MEG data to 0.1–30 Hz for visualization of the evoked responses and for sensor- and source-level decoding. Ocular artifacts were suppressed by removing those independent components (1–4 per subject, on average 3) that correlated most with the EOG signal.

For the subsequent data analysis, only planar gradiometers were used due to the straightforward interpretation of their spatial pattern; they show the maximum signal right above the active source.

Epochs with two different pre- (200 ms and 500 ms) and post-stimulus (800 ms and 1500 ms) periods were extracted from the MEG data at every word stimulus. Epochs were rejected if any of the planar gradiometer signals exceeded 4000 fT/cm. Deviant epochs were excluded from data analysis. Fig. 6C illustrates which evoked responses to what stimuli were employed in decoding attention direction.

The trial counts were equalized, and the responses averaged across each condition (“Attended Yes”, “Attended No”, “Unattended Yes” and “Unattended No”) for visualization and source estimation.

### Source estimation

Head models were constructed based on individual magnetic resonance images (MRIs) by applying the watershed algorithm implemented in the FreeSurfer software [46,47, *Version* 5.3;]. Using the MNE software, single-compartment boundary element models (BEM) comprising 5120 triangles were then created based on the inner skull surface. In addition to the one subject with technical problems in MEG recording, the MRIs of three subjects were not available, leaving 11 subjects for the source estimation.

For the source space, the cortical mantle was segmented from MRIs using FreeSurfer and the resulting triangle mesh was subdivided to 4098 sources per hemisphere. The dynamic statistical parametric mapping [48, *dSPM*;] variant of minimum-norm estimation was applied to model the activity at these sources. The noise covariance was estimated from the 2-min resting-state measurement of each subject. These data were pre-processed similarly as the task-related data.

The source amplitudes for the attention conditions “Attended Yes”, “Unattended Yes”,”Attended No” and “Unattended No” were estimated for all subjects individually. For the group-level source estimate, the obtained source amplitudes were first normalized such that the absolute peak value of the attended condition became one, the estimates were morphed to the FreeSurfer average brain and then averaged across subjects.

### Decoding

#### Sensor-level decoding

A linear support vector machine [49, *SVM*;] classifier implemented in the Scikit-learn package^45^ was applied to unaveraged epochs to decode the conditions “Attended Yes” vs. “Unattended Yes” and “Attended No” vs. “Unattended No”. For comparison, decoding was also performed stimulus-word-wise, i.e. “Attended Yes”+“Unattended Yes” vs. “Attended No”+“Unattended No”. In addition, pitch-wise and single pitch variant attention-wise decoding were performed.

The pre-processed MEG data (filtered to 0.1–30 Hz) were down-sampled by a factor of 8 to a sampling rate of 125 Hz to reduce the number of features while preserving sufficient temporal information. Amplitudes of the planar gradiometer channels were concatenated to form the feature vector. Shuffled five-fold cross-validation (CV) was applied with an 80/20 split; 80% of data were used for training and the rest for testing. The empirical chance level was around 55% for our sample size of 500 epochs in this two-class decoding task^50^. We also verified the empirical chance level by the method by Ojala and Garriga^51^ and in our case it was 53%.

Decoding was separately performed on data of 1) the entire 2-s epoch (250 time points x 204 channels; *long-epoch decoding*) 2) the entire 1-s epoch (125 time points x 204 channels; *short-epoch decoding*), 3) one time point (1 time point x 204 channels; *time-resolved decoding*), and 4) one channel (250 time points x 1 channel; *spatially-resolved decoding*).

#### Source-level decoding

A linear SVM decoder with five-fold cross-validation (80%/20% split for training/testing) was applied to the individual source estimates for the attention conditions “Attended Yes” vs “Unattended Yes” and “Attended No” vs “Unattended No” calculated from all MEG planar gradiometer channels. A spatial searchlight decoding across the source space was used on the 1-s (−200–800 ms after stimulus onset) epochs, and the resulting accuracy maps were morphed to the FreeSurfer average brain and averaged across the subjects (N = 11). The accuracy maps for attention conditions “Attended Yes” vs “Unattended Yes” and “Attended No” vs “Unattended No” were then averaged to obtain a general accuracy map.

## Data availability

The datasets generated and analysed during the current study are not publicly available due to the local legislation on research on humans but are available from the corresponding author on reasonable request.

## Acknowledgements

This research was supported by Academy of Finland, grant No. 295075 “NeuroFeed”, and European Research Council, grant No. 678578 “HRMEG”. The content is solely the responsibility of the authors and does not necessarily represent the official views of the funding organizations. The measurements were conducted at the MEG Core of Aalto Neuroimaging Infrastructure, Aalto University, Finland, and financially supported by Aalto Brain Centre. The authors thank prof. Ville Pulkki and Aalto Acoustics Lab, Aalto University, for the loan and guidance on the use of the dummy head.

## Author contributions

D.K., H.K., M.J., A.V. and L.P. designed the experiment. D.K., H.K. and M.J. implemented the experiment. D.K. and H.K. conducted the measurements. D.K. analysed the results and wrote the manuscript. All authors reviewed the manuscript.

